# Human genetic studies and zebrafish models identify Plxna4 as a regulator of adiposity, somatic growth, and feeding behaviours

**DOI:** 10.1101/2025.03.15.643290

**Authors:** Panna Tandon, Zinnia Lyall, Madeleine Cowie, James E. N. Minchin

## Abstract

Obesity is a major public health crisis, affecting billions worldwide and increasing the risk of metabolic and cardiovascular diseases. While lifestyle factors play a role, genetic variation is a key determinant of both obesity susceptibility and the efficacy of treatment strategies. Recent studies have implicated the Semaphorin 3 signalling pathway in obesity; however, specific roles for pathway components remain largely unexplored. Here, we focus on Class A Plexins and their potential contributions to body weight regulation. Using large-scale genetic association data, we identified that rare, predicted loss-of-function mutations in *PLXNA4* were associated with body mass index (BMI) in females. Furthermore, common variant analysis revealed that genetic variation at *PLXNA4* was linked to BMI, height, and various neuropsychiatric disorders. To investigate the biological role of Plxna4, we generated zebrafish *plxna4* loss-of-function mutants, which exhibited an 85–92% reduction in Plxna4 protein. Despite appearing morphologically normal, mutant zebrafish at juvenile stages were shorter, had increased body fat levels relative to size-matched wild-type siblings, and displayed hypertrophic subcutaneous adipose tissue. Feeding assays revealed that *plxna4* mutants consumed more food than wild-type siblings and exhibited food-stimulated hyperactivity, characterised by increased swimming speed, higher speed variability, and frequent high-speed bursts. Together, these findings demonstrate a conserved role for Plxna4 in regulating feeding behaviour and body fat levels, providing new insights into the genetic basis of obesity and warranting further studies to elucidate the molecular mechanisms underlying these effects.

## Introduction

Obesity, defined by body mass index (BMI), constitutes an increasingly significant public health concern, with rates of adult obesity doubling since 1990 and adolescent obesity quadrupling (1). Obesity reduces life expectancy and is a risk factor for numerous chronic diseases, including cancers, cardiovascular disorders, and diabetes (2). Given its profound impact on global health, understanding the causes of obesity and factors that predict the risk of development remains a critical objective. Although the aetiology of this condition is multifactorial, research has supported a strong underlying genetic component (3). For instance, Maes et al. (1997) reported that genetics account for 50% to 90% of the variance in BMI and other adiposity measures (4). Genetic factors also seem to influence an individual’s response to obesity therapeutics (5). Thus, investigating the genetic variation that influence susceptibility to obesity offers a promising avenue for identifying vulnerable individuals and developing patient-individualised therapeutic interventions.

The Semaphorin 3 signalling pathway is comprised of three key components: extracellular Semaphorin 3 signalling proteins (SEMA3s), which interact with Class-A Plexin receptors (PLXNAs) and Neuropilin co-receptors (NRPs) (6). This pathway plays a crucial role in neuronal cytoskeletal remodelling, influencing axonal development, circuit formation, synapse formation, axonal pruning, and axon terminal branching (6). In individuals with severe early-onset obesity, van der Klaauw et al. (2019) identified 40 rare variants in genes belonging to the Semaphorin 3 signalling pathway. Moreover, variants within this pathway were enriched within 982 severely obese individuals compared to controls (7). Follow-up mechanistic analyses demonstrated that Semaphorin 3 signalling influences the development of hypothalamic melanocortin circuits, which are essential for regulating feeding behaviours, energy homeostasis and obesity susceptibility. However, the precise roles of individual pathway components in obesity remains unclear. Notably, rare obesity-linked variants within the four *PLXNA* genes (*PLXNA1-4*) were shown to affect cell surface localisation and cellular collapse in vitro (7). Despite these findings, the functional roles and mechanisms of *PLXNA* genes in energy homeostasis are lacking. Intriguingly, a recent case report linked a *PLXNA4* variant to exaggerated weight loss outcomes (8), suggesting Plxna4 could provide insights into personalised obesity treatments.

Zebrafish serve as an excellent model for studying energy homeostasis due to their highly conserved hypothalamic circuitry, which regulates food intake, energy balance, somatic growth, and body fat accumulation. The Pomc and Agrp neurons in zebrafish are located within hypothalamic regions with similar functionality to the mammalian arcuate nucleus (9), and their melanocortin system plays a central role in regulating somatic growth. Specifically, Agrp neurons positively reinforce growth hormone (GH)-induced growth and body fat accumulation (9,10). Whereas Pomc neurons repress GH-induced growth (11). Supporting this, *gh1* mutant zebrafish exhibit short stature and extreme adiposity, mirroring aspects of human growth hormone deficiencies (12). In addition to regulating growth, zebrafish Pomc neurons have long-range projections to Mc4r-expressing neurons in the spinal cord, which modulate swimming and feeding behaviours (13). Whether Plxna4 regulates these circuits in currently unknown. Recent studies have also highlighted the caudal and lateral hypothalamic units as key regulators of appetite control in zebrafish (14,15). These regions exhibit anti-correlated activity patterns during hunger and feeding states, indicating specialised roles in appetite regulation (14). The lateral hypothalamus is particularly active during feeding and promotes food-seeking behaviours, mirroring its mammalian counterpart’s role in appetite-driven actions (14,15). These findings reinforce the zebrafish as a powerful system for dissecting the neural mechanisms of energy balance, with potential implications for understanding human metabolic disorders.

In this study, we combined human genetic association analyses with functional studies in zebrafish to explore a role for Plxna4 in feeding and obesity susceptibility. We first examined rare and common *PLXNA4* variants associated with BMI and related traits, then generated *plxna4* loss-of-function zebrafish mutants to assess their metabolic and behavioural phenotypes. Our results reveal that *plxna4* mutants exhibit increased body fat, hyperphagia, and food-stimulated hyperactivity, supporting a conserved role for Plxna4 in energy balance.

## Methods

### Rare variant associations linking BMI and class-A Plexins

Exome-based association statistics from the UK Biobank were accessed using the Genebass resource (16). We focussed on predicted loss-of-function (pLOF) variants as they are most likely to have a strong impact on gene function. BMI in three cohorts (male, female, and combined) was examined to account for potential sex-specific genetic effects. Pre-computed SKAT-O tests were used to assess mutational burden as single variants often had directionally opposite effects (**Table S1**). FDR (Benjamini-Hochberg) correction was applied to the p-values across all tests (12 tests performed in total, 1 variant category × 1 statistical test x 3 traits x 4 genes, significance threshold was p = 0.00417). GnomAD v4.1 was used to identify pLOF variants in *PLXNA4* (17), and LOEUF score was used to assess observed/expected frequency of pLOF variants in *PLXNA4*.

### *PLXNA4* genome-wide association data, eQTLs and open chromatin landscapes

Published GWAS summary statistics for *PLXNA4* were downloaded from the GWAS Catalog (18). 78 associations were identified and 49 that were genome-wide significant (p < 5×10−8). The LDproxy function within the LDLinkR R package was used to identify proxy SNPs in high linkage disequilibrium with the GWAS SNP (>0.7 R2), in the CEU European population (GRCh38 high coverage) and within a 1Mb window. Pre-computed significant single-tissue eQTLs for *PLXNA4* were downloaded from the GTEx portal v10 (19). Overlap between GWAS variants and eQTLs was performed using the GenomicRanges package in R (20). GWAS variant and eQTL colocalisation analysis was performed using the eQTpLot R package (21). Open chromatin regions from ATAC-Seq within arcuate-like hypothalamic neurons and their progenitors (derived from pluripotent embryonic stem cells) were obtained from (22). Micro-C data from H1-hESCs were obtained to assess chromatin interactions (23).

### Zebrafish husbandry and maintenance

Zebrafish were housed within the Queen’s Medical Research Institute at the University of Edinburgh on a recirculating system with a 14h:10h light:dark cycle. From 5 days post fertilisation (dpf), larvae were raised in static water with slow drip water flow beginning from 10 dpf. Fish were raised at ∼8-9 fish per L. From 5-30 dpf, fish were fed four-times daily with two live feeds (*Artemia salina* and *Brachionus plicatilis* L-Strain rotifer from Zebrafish Management Ltd) and twice daily with a dry feed (ZM100-400 and GM micro 75-500). From 30 dpf, fish were fed three-times daily with one live feed (*Artemia salina*) and two dry feeds (ZM100-400 and GM micro 75-500). The first feed was given between 9-10am and the last feed between 3-4pm. All zebrafish experiments were authorised under the Animals (Scientific Procedures) Act 1986 (project licence PP9112175).

### CRISPR mutagenesis and genotyping

CRISPR mutagenesis of zebrafish embryos was conducted using standard procedures (24). Briefly, guide RNAs (gRNAs) to target zebrafish *plxna4* were designed using UCSC Genome browser track ZebrafishGenomics (Varshney et al; 2015). gRNA targets were selected based on off-target scores <250. CHOPCHOP software was later used to verify potential off-target and efficiency scores. Primers (Integrated DNA Technologies) were ordered and annealed to make the gRNA template for synthesis using an mMessage mMachine in vitro SP6 transcription kit (Invitrogen, Cat. number AM1340). RNA was cleaned using zymoclean column kits (Zymo Research, Cat. number R1013) and quantified using a Nanodrop spectrophotometer. gRNA was incubated with Cas9 protein (New England Biolabs, Cat. number M0646T) and heated for 5 mins at 37°C, for a final concentration of 133 pg/nl gRNA and 266 pg/nl Cas9 protein. Embryos were injected with 1 nl at the one-cell stage and raised for genotyping using standard PCR amplicon T7E1 assay (New England Biolabs, Cat. number M0302S) to confirm genome editing, grown to adulthood and outcrossed to identify and generate stable lines. PCR amplicons from the *plxna4* locus were cloned (Agilent Strataclone PCR cloning kit, Cat. Number 240205) and sequenced to determine the *plxna4*^*ed133*^ allele. Subsequent genotyping of *plxna4*^*ed133*^ larvae was conducted on whole-larvae or tail clip genomic DNA using GoTaq (Promega) standard PCR conditions utilising a three-primer strategy to identify allele-specific amplicons and the wildtype, heterozygous and homozygous genotypes. Kompetitive Allele Specific PCR (KASP) on Demand genotyping kit with *plxna4*^*ed133*^ allele-specific probes was also used for genotyping following manufacturer’s protocols (LGC Biosearch Technologies, project number 2044_005). Primers were: *plxna4* CRISPR gRNA target sequence: GAATACAGTACGTTTCGGGTGG. *plxna4*^*ed133*^ genotyping primers: forward – 5’ TTTCTTGCTGCGTTGTTGTC, reverse -5’ TAACGCAACCAAGAGGAACC. *plxna4*+ allele-specific genotyping primer: reverse – 5’ TTTTCCACCCGAAACGTACT. *plxna4*^*ed133*^ allele-specific genotyping primer: reverse – 5’ GTTTTCCACACAGAATGTAACG.

### Western blotting

Western blotting was performed on dissected wild-type or *plxna4*^*ed133*^ adult male zebrafish brains to assess Plxna4 protein abundance. Briefly, brains were dissected and placed on ice in PBS before being dissociated in RIPA buffer (Pierce) with protease inhibitors. Tissue was disrupted using pestle plunges as well as 5 × 30 second rounds of water-bath sonication (Diagenode). Lysate was centrifuged and quantified using standard BCA assay (23225, Thermo Fisher Scientific) and Tecan plate reader. Protein lysate was boiled at 95°C for 10 minutes together with loading dye and 50 ug loaded on each lane of a 8% Bis-Tris gel with 1 x MOPs (Invitrogen) and transferred at 10V overnight at 4°C. Standard antibody stains were performed with 5% milk/TBST blocking solution. Primary antibodies were rabbit anti Plxna4 (Abcam ab39350) and rabbit anti-ERK (Cell Signalling Technology 1:2000). LiCOR secondary antibodies were mouse 680 and rabbit 800 1:10,000. Protein expression was imaged using a LiCOR Odyssey CX and quantified using LiCOR Empiria Studio Software against total protein REVERT700 stain and found a 6.5-fold reduction (85% loss). This measurement was replicated using ImageJ quantification and with total ERK as a loading control and found 12.8-fold reduction (92.2% reduction) in *plxna4*^*ed133*^ mutants.

### Food intake and locomotion assays

Voluntary food intake was assessed in 30 dpf zebrafish, which were fasted for 12h prior to moving individual wild-type or *plxna4* mutant fish to single wells of a six-well plate. Defined amounts of a live feed (*Artemia*) were added to each well and the number of *Artemia* ingested within a 10-minute period was recorded. Intake differences were assessed using a *X*^*2*^ test. The same feeding assay was performed with the addition of video recording of fish locomotion over a 10-minute period at 0.07 s/frame.

### Nile Red staining and high-fat diet manipulation

Fish were incubated in Nile Red (Thermo Fisher, Cat number N1142) according to previous protocols (25). Nile Red stained fish were anaesthetised in MS222, mounted in 3% methylcellulose and imaged on a Leica M205 stereomicroscope. Nile Red images were initially processed in FIJI/ImageJ, and segmentation was performed in Ilastik software (v1.4.0) using a Pixel Classification segmentation workflow. Segmented masks were imported back to FIJI/ImageJ for quantification. We used Nile Red to also assess lipid droplet (LD) morphology, and imaging was performed on a Zeiss LSM710 confocal microscope. For the high-fat diet assay, juvenile fish from a wildtype and *plxna4*^*ed133/ed133*^ incross were genotyped at 27 dpf, and size matched (mean 8,889 μm and 8,426 μm for each group respectively) and distributed into high-fat diet or control fed groups (n=8 each). Pairs of fish were placed into mesh containers within a 10 L system tank for the duration of the experiment and fed according to standard husbandry conditions by facility staff. Container positions were rotated randomly daily during the assay to minimise position effects. Fish were taken off system and placed into 200 ml system water (control) or 5% chicken egg yolk (Sigma, Cat. number E0625) for two hours daily in the morning for 14 days (26). Fish were incubated in neutral lipid dye Nile Red before and after diet treatment as above.

### Statistical analysis

Linear mixed-effects models were used to compare standard length, adiposity (Nile Red area) and lipid droplet diameters between mutant and wild-type groups, accounting for variability across 8 independent experiments. Mixed-effects ANCOVA was used to assess differences in square root-transformed Nile Red area between wild-types and mutants, while controlling for standard length as a covariate and accounting for batch variability across experiments. A two-way ANOVA was used to test for locomotion response to diet (pre-food and post-food). Behavioural responses to food in wild-types and mutants were compared using Welch two-sample t tests.

## Results

### Rare predicted loss-of-function variants in *PLXNA4* are associated with BMI in females

We analysed publicly available exome sequencing data for rare variant associations between class-A Plexins and BMI. We assessed gene-level associations of pLOF variants with BMI in three cohorts: females (n = 213,209), males (n = 180,591), and the combined population (n = 393,562) (**Table 1**). Across all four *PLXNA* genes, statistical associations were weak, possibly due to the rarity of these variants and/or small effect sizes. However, after correcting for multiple tests, mutational burden in *PLXNA4* remained significantly associated with BMI in females (**Table 1**). Individual variants exhibited both positive and negative effects on BMI, but none reached statistical significance (**Table S1**). In addition to the four coding variants identified previously (7), 105 *PLXNA4* pLOF variants were identified in gnomAD v4.1, with a LOEUF score of 0.22 indicating strong negative selection against PLXNA4 loss-of-function mutations (**Fig. 1A & Table S2 & S3**). Notably, pLOF variants were more frequent at the junction between the Sema and PSI domains, suggesting this region is under reduced evolutionary constraint (**Fig. 1B**). In summary, rare predicted loss-of-function variants in *PLXNA4* show a significant association with BMI in females, suggesting a sex-specific genetic influence for Plxna4 on body mass regulation.

**Table 1:** Gene burden association statistics for BMI traits in Class-A Plexins 1-4. Significant association marked by asterisk after adjusting for multiple tests (significance threshold, p = 0.00417).

| Gene | Trait | P-value (SKAT-O) | Adjusted P-value (SKAT-O) | pLOF variants | Cumulative allele frequency (pLOF) |
| --- | --- | --- | --- | --- | --- |
| *PLXNA1* | BMI (combined) | 0.344 | 0.688 | 17 | 3.17e-5 |
|  | BMI (female) | 0.229 | 0.588 | 11 |  |
|  | BMI (male) | 0.913 | 0.928 | 9 |  |
| *PLXNA2* | BMI (combined) | 0.928 | 0.928 | 76 | 1.9e-4 |
|  | BMI (female) | 0.614 | 0.819 | 47 |  |
|  | BMI (male) | 0.867 | 0.928 | 48 |  |
| *PLXNA3* | BMI (combined) | 0.49 | 0.735 | 81 | 2.49e-4 |
|  | BMI (female) | 0.474 | 0.735 | 61 |  |
|  | BMI (male) | 0.0632 | 0.253 | 33 |  |
| *PLXNA4* | BMI (combined) | 0.0171 | 0.103 | 36 | 5.19e-5 |
|  | BMI (female) | 0.00324* | 0.039* | 17 |  |
|  | BMI (male) | 0.245 | 0.588 | 21 |  |

**Figure 1.**
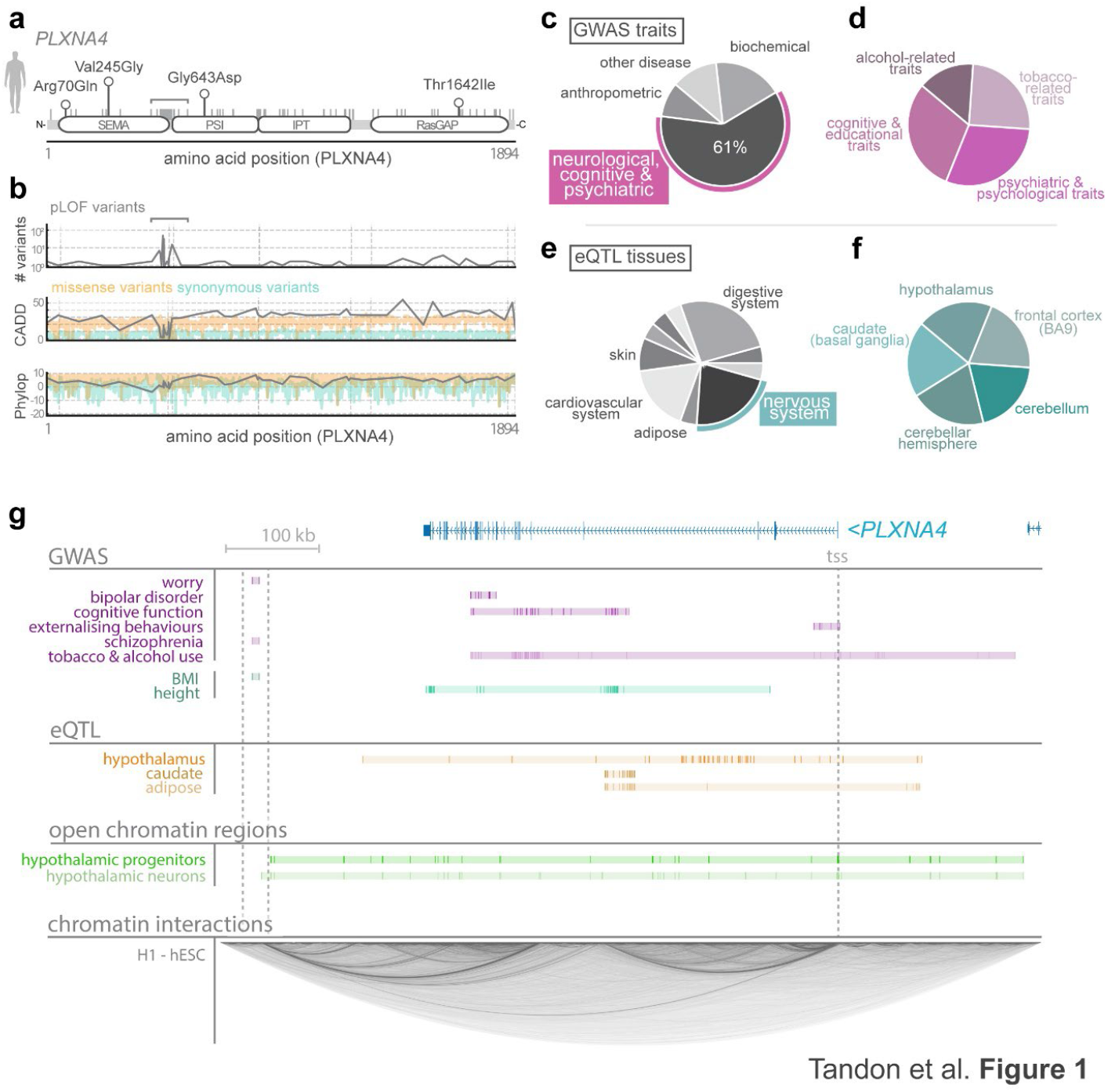
Human genetic data linking *PLXNA4* to BMI, height and a range of neuropsychiatric disorders. **(A)** Schematic showing the domain structure of human Plxna4, including Semaphorin domain (SEMA), PSI and IPT domains, and RasGAP domain. Grey vertical lines indicate location of pLOF variants within Plxna4 from gnomAD database. Four obesity-linked variants from van der Klaauw are highlighted with lollipops. Amino acid position is indicated on the x-axis. Bar indicates permissive region for variants. **(B)** Graphs showing number of pLOF variants at each position of the PLXNA4 gene (upper graph), CADD scores for pLOF variants (grey), missense (yellow) and synonymous (green) variants (middle graph) and Phylop scores for pLOF (grey), missense (yellow) and synonymous (green) variants (lower graph). Amino acid position is shown on x-axis. **(C)** 49 GWAS SNPs at PLXNA4 grouped according to trait. 61% of GWAS SNPs at PLXNA4 are associated with neurological, cognitive and psychiatric traits. **(D)** Subdivision of the 61% GWAS SNPs from C in to groups including tobacco-related traits use, alcohol-related traits, cognitive and educational traits and psychiatric and psychological traits. **(E)** Pie chart to show the tissues with significant PLXNA4 eQTLS based on GTEx data. **(F)** Subdivision of nervous system eQTLs show PLXNA4 eQTLs in hypothalamus, cerebellum, caudate and frontal cortex. **(G)** Locus view of human PLXNA4 highlighting location of GWAS SNPs, tissue-specific eQTL, open chromatin regions in hypothalamic neural progenitors and differentiated neurons, and chromatin interactions.

### Common variants at *PLXNA4* are associated with BMI, height and a broad range of neurological and psychiatric disorders

Focussing on *PLXNA4*, we leveraged published genome-wide association studies (GWAS) data to examine common-variant associations. Using a genome-wide significance threshold of p<5×10−8, we identified 49 associations between variants at the *PLXNA4* locus and a broad range of traits (**Fig. 1C & Table S4**). Notably, genetic variants at *PLXNA4* were associated with BMI and height (**Fig. 1C**). However, strikingly, 61% of GWAS variants at this locus were linked to neurological, cognitive and psychiatric traits-including worry, bipolar disorder, cognitive function, externalising behaviours, schizophrenia, and alcohol and tobacco use suggesting pleiotropic roles for Plxna4 in weight regulation, somatic growth and neuropsychiatric disorders (**Fig. 1C & D**). Single-tissue eQTLs for *PLXNA4* were identified in both brain and peripheral tissues, including the hypothalamus, caudate (basal ganglia), cerebellum, and frontal cortex – areas with known roles in feeding and obesity (**Fig. 1E & F**). Of the 49 GWAS SNPs at *PLXNA4*, 9 were also associated with *PLXNA4* mRNA expression within multiple tissues, including the cerebellum (**Table S5**), and 2.7% and 0.4% of GWAS variants (and their proxy SNPs) directly overlapped with open chromatin regions in hypothalamic neuron progenitors or differentiated neurons respectively (**Table S6**). Whilst the BMI-associated variants were located downstream of *PLXNA4*, extensive chromatin interactions were evident between this region and other sites across the locus including to the *PLXNA4* transcriptional start site (**Fig. 1G**). Altogether, GWAS data reveals common variants at *PLXNA4* that are associated with BMI, height, and a range of neuropsychiatric traits.

### Zebrafish *plxna4* mutants are smaller than wild-types but have increased body fat

To understand Plxna4 roles in body weight regulation, height and neuropsychiatric traits we turned to the zebrafish model system. Zebrafish Plxna4 is highly conserved with its mammalian counterparts, both in sequence identity and syntenic organization (**Fig. S1A & B**). Using established CRISPR methodology, we generated a new *plxna4* allele (ed133) containing a Phenylalanine-to-Leucine change at amino acid 44, followed by a frameshift, a nonsense 29 amino acid sequence and a premature stop codon (**Fig. 2A**). By Western blot, we observed an 85-92.2% reduction in Plxna4 protein, suggesting ed133 is a strong loss-of-function allele (**Figs. 2B & S2**). In accordance with a previous study, we observed no gross morphological or mortality issues in *plxna4*^*ed133*^ mutants (not shown) (27). However, in juvenile animals between 30-40 dpf, *plxna4*^*ed133*^ homozygous mutants were significantly shorter than wild-type siblings (p < 0.0001) (**Fig. 2C & D**). Without accounting for animal size, the *plxna4* mutants had significantly lower adiposity relative to wild-type siblings (p = 0.0021, not shown). However, when size was controlled for, *plxna4* mutants had significantly higher adiposity levels than size-matched wild-type siblings (p = 0.005), suggesting that *plxna4* mutants accumulate greater body fat than wild-types (**Fig. 2C & F**). High-resolution imaging of lipid droplets (LDs) within subcutaneous adipose revealed *plxna4* mutants had larger LDs than wild-type siblings, suggesting an overall hypertrophic morphology in subcutaneous adipose (p < 0.001) (**Fig. 2G**).

**Figure 2.**
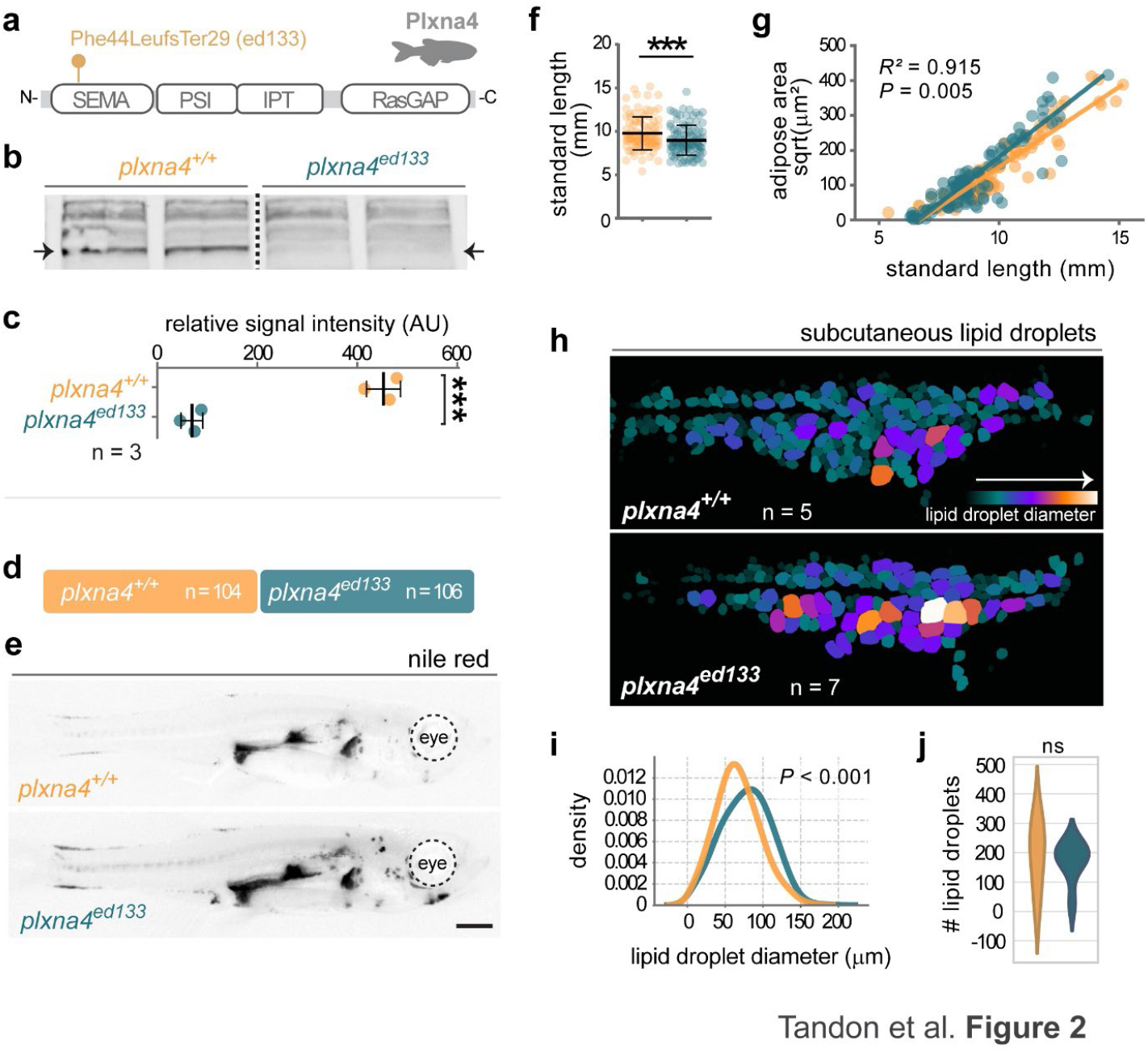
Zebrafish loss-of-function *plxna4* mutants are shorter and have increased adiposity and hypertrophic subcutaneous adipose tissue. **(A)** Schematic of zebrafish Plxna4 showing the location of the ed133 mutation within the Semaphorin-binding domain. **(B)** Western blot showing the absence of Plxna4 in the ed133 homozygous zebrafish adult brains. Arrows denote location of Plxna4 band. **(C)** Quantification of protein loss form Western blots revealed 85-92% reduction in Plxna4 protein in ed133 homozygotes. **(D)** Nile Red data consisted of 210 observations from 8 independent experiments, with 106 samples from the mutant group and 104 samples from the wild-type group. **(E)** Nile Red images showing increased body fat levels in plxna4 ed133 homozygotes. **(F)** A linear mixed-effects model was used to compare standard length between mutant and wild-type groups, accounting for variability across 8 independent experiments. **(G)** Linear mixed-effects ANCOVA model of square root-transformed Nile Red area as a function of standard length and group (mutants vs. wild-types), with experiment as a random effect to account for experimental batch effects. Solid lines indicate group-specific regression lines. The model explained 91.5% of the variance in √Area (Conditional R^2^). **(H)** Segmented lipid droplets from the lateral subcutaneous adipose tissue (LSAT), colour-coded according to diameter. **(I)** Probability density function of lipid droplet diameter by group. A linear mixed-effect model was used to account for batch effects and variability between experiments. **(J)** The number of lipid droplets was not significantly different between groups.

### Zebrafish *plxna4* mutants are hyperphagic

Appetite and food consumption play central roles in the development of obesity, with the majority of known genetic contributions to human obesity altering appetite regulation and food intake. Given this, we next investigated whether food intake was altered in *plxna4* zebrafish mutants. We tested voluntary food intake in *plxna4* mutants at 30 dpf. Following an initial 12 h fasting period, we provided individual fish with a defined amount of a live feed (*Artemia*) and recorded the number of *Artemia* ingested over a 10-minute period (**Fig. 3A**). Over the 10 min period, wild-type zebrafish exhibited a broad range of food intake levels, with the majority consuming less than 75% of the available food (**Fig. 3B**). In contrast, the majority of *plxna4* mutants consumed more than 75%, indicating a shift toward increased food intake in mutant fish (p = 0.012) (**Fig. 3B**). We also performed a 14-day high-fat diet exposure shown previously to increase adiposity and circulating lipid (**Fig. S3**) (26). Following 14-days of the high-fat diet, both wild-type siblings and *plxna4* mutants had increased adiposity; however, the rate of adipose increase was similar across both genotypes, suggesting a similar capacity to accumulate lipid in adipose after a high-fat diet (**Fig. S3**).

**Figure 3.**
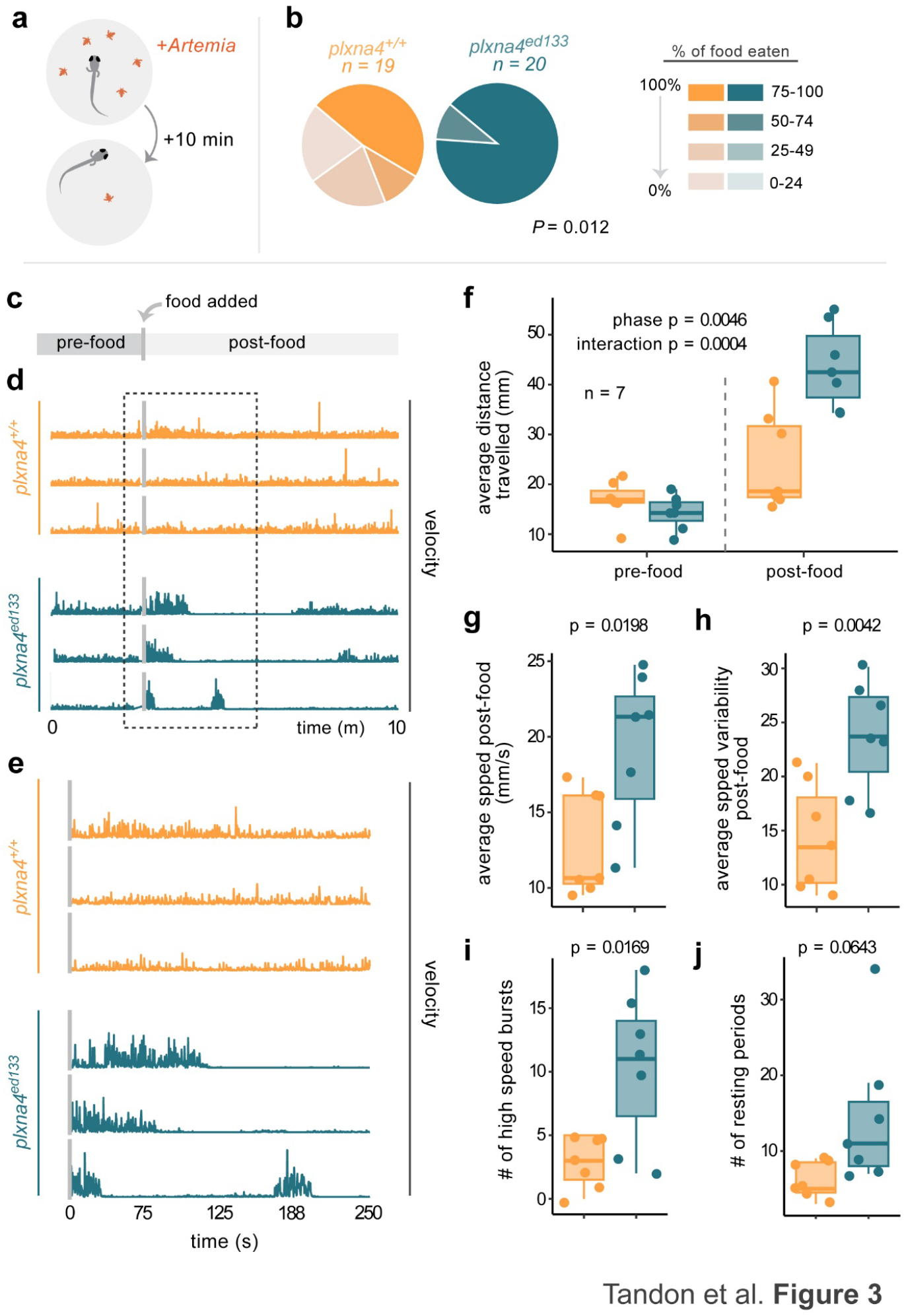
Zebrafish *plxna4* mutants are hyperphagic and show food-stimulated hyperactivity. **(A)** Schematic showing food (*Artemia*) intake assay over 10-minute duration. Fish were 30 dpf. **(B)** Quantification of food intake after a 10-min period. Intake levels grouped into quartiles and a Chi-squared statistical test. Increased levels of food intake are shown in stronger colours. **(C)** Schematic showing locomotion assays with pre-food and post-food phases. *Artemia* was added at the grey vertical line. Fish were 30 dpf. **(D)** Velocity tracks over the course of the locomotion assay are shown for three representative wild-type and *plxna4* mutants fish. Dotted box represents time period shown in E. **(E)** Zoomed in area immediately following addition of food. **(F)** Two-way ANOVA showing effect of phase (pre-food or post-food) and group (mutant or wild-type) on average distance travelled. Phase and the interaction between phase and group/genotype in significant. **(G)** Average speed in post-food phase was statistically increased in *plxna4* mutants. **(H)** Average speed variability was statistically increased in post-food phase in mutants. **(I)** The number of high-speed bursts was statistically higher for *plxna4* mutants in the post-food phase. **(J)** Number of resting periods was higher in *plxna4* mutants but did not reach statistical significance.

**Figure 4.**
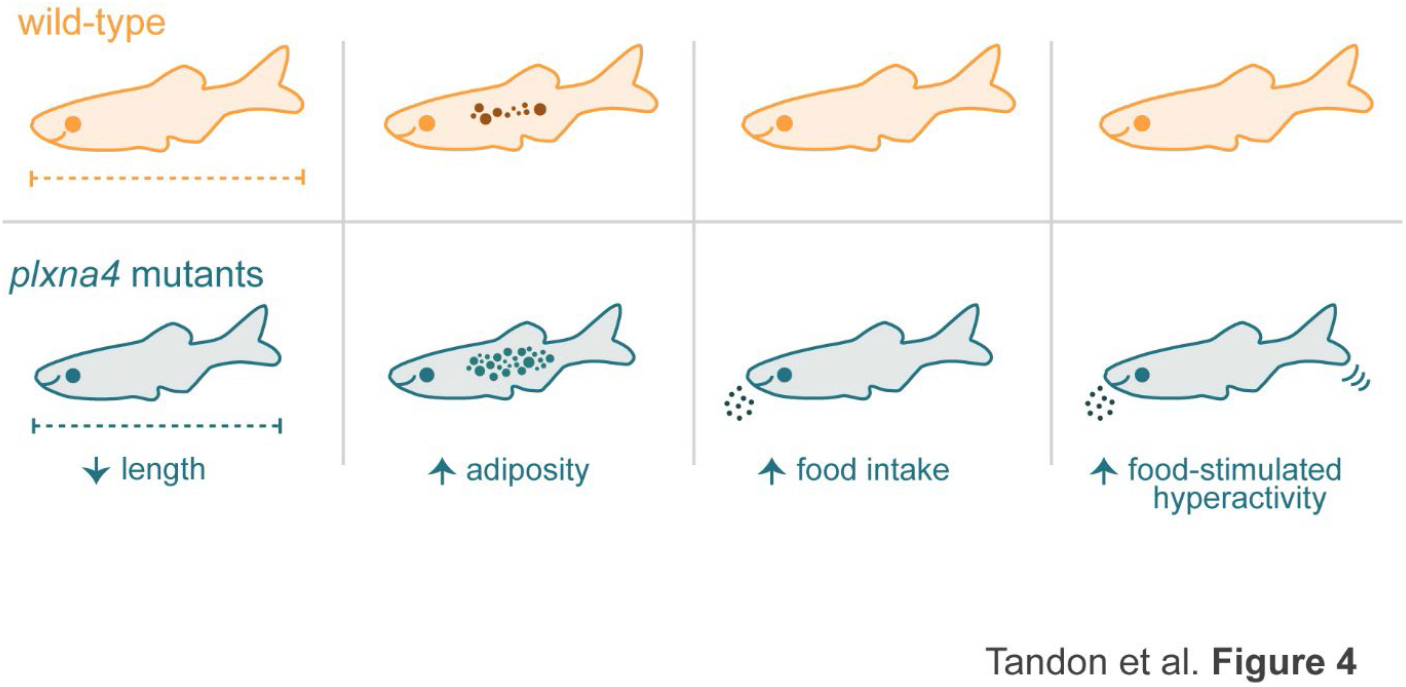
Overview of baseline *plxna4* mutant phenotypes in zebrafish. Compared to wild-type siblings, *plxna4* ed133 mutants have a reduced standard length, have increased body fat levels/adiposity, show increased food intake and have a series of food-stimulated hyperactivity behaviours including increased average speed in response to food, increased speed variability in response to food and increased high-speed swimming bursts in response to food.

### Mutation of *plxna4* in zebrafish results in feeding-related hyperactivity

Feeding is governed by structured, coordinated sequences of behaviours that regulate food intake. Therefore, we next assessed behavioural responses to food in *plxna4* mutants. Behavioural assays were conducted in 30 dpf zebrafish over 10 minutes (**Fig. 3C**). At baseline, before *Artemia* were added to the environment, *plxna4* mutants had a slightly lower average distance travelled relative to wild-type controls, although this wasn’t statistically significant (**Fig. 3F**). However, following addition of food, *plxna4* mutants showed an increase in average distance travelled, indicating a greater activity level in response to food in mutants (p = 0.0004) (**Fig. 3F**). The greater food-stimulated activity in *plxna4* mutants was characterised by a higher average speed (**Fig. 3G**), greater speed variability suggesting more erratic movements (**Fig. 3H**), and an increase in high-speed burst swimming (**Fig. 3I**). *plxna4* mutants also had greater rest period, although this was not statistically significant (**Fig. 3J**). Overall, these data show that in response to food *plxna4* mutants have hyperactive periods characterised by greater speeds and speed variability.

## Discussion

We leveraged publicly available rare and common genetic association data to review links between Plxna4 and body weight regulation. Focusing on rare, coding variants, we used pre-computed association statistics from the Genebass resource and identified that mutational burden of pLOF variants in *PLXNA4* were associated with BMI in females, suggesting a potential sex-specific role for Plxna4 in weight regulation. The reason for this female-specific association is unclear; however there are well characterised body fat differences between males and females, including higher overall body fat levels in females, distinct fat distributions between the sexes, and varying disease risks at equivalent BMIs between sexes (28). Notably, rare variants within multiple genes, including *DIDO1, PTPRG*, and *SLC12A5*, also showed female-specific associations with BMI at exome-wide significance, supporting the idea of sex-specific genetic regulation of body weight (29). Based on these data, it will be important to assess sex-specific roles for Plxna4 in model systems. We acknowledge that the association between variants in *PLXNA4* and BMI was not statistically significant at an exome-wide level; however, common variant association data from GWAS also identified associations with BMI at the *PLXNA4* locus, along with height and a range of neuropsychiatric traits. These findings suggest pleiotropic roles for Plxna4 in the brain influencing appetite, feeding behaviour and BMI, alongside neuropsychiatric traits. Prior studies have highlighted genetic overlap between BMI and psychiatric disorders, raising the possibility that these traits share common neurobiological mechanisms or inter-dependence (30,31). Further investigation is needed to dissect these relationships and roles for Plxna4 in model systems.

To determine the contribution of Plxna4 to weight regulation, we generated new loss-of-function *plxna4* zebrafish mutants. The *plxna4* mutants exhibited both reduced body length and increased body fat levels – both traits identified by GWAS - suggesting they provide a valuable model to explore Plxna4 roles in human disease processes. Western blot analysis confirmed an 85-92% reduction in Plxna4 protein, indicating that the mutation represents a strong loss-of-function allele. A recent study characterised *Plxna4* null mice and reported that mutants were smaller than wild-type control animals and resistant to high-fat diet induced obesity (32). We also observe that zebrafish *plxna4* mutants are smaller and administering a high-fat diet to the mutant zebrafish did not result in exacerbated adipose accumulation, however the high-fat diet was only administered for 14 days, and our sample size was low suggesting caution in interpretation. Further, *plxna4* mutants initially appear to have reduced adiposity if only absolute adipose size is considered. However, once body size was accounted for, the zebrafish mutants had significantly higher relative body fat levels. Given our large sample sizes (>100 zebrafish) we are able to detect relatively small effect sizes, which may be important for studying these adiposity phenotypes.

Intriguingly, despite increased food intake, *plxna4* mutants exhibited reduced standard length suggesting additional underlying factors influence their growth. Interestingly zebrafish growth hormone (*gh1*) mutants also display increased adiposity alongside reduced somatic growth (12), raising the possibility that Plxna4 deficiency may impair GH signalling. Notably, obesity and hyperinsulinemia can blunt the GH axis (33), potentially contributing to the growth phenotype observed in *plxna4* mutants. Notably, in zebrafish Pomc neurons were found to inhibit GH production via Somatostatin neurons (11). It will be interesting to study whether Plxna4 influences wider patterning of Pomc neurons, which show large functional and molecular heterogeneity to regulate a diverse range of processes involved in energy homeostasis (34–37). Alternatively, broader hypothalamic dysfunction in *plxna4* mutants could underlie their short stature as GH, thyroid hormones, and adrenal hormones all play critical roles in somatic growth (38). Additionally, Plxna4 may have a direct role in skeletogenesis, myogenesis or exert broader effects on metabolism. Future studies should explore these potential mechanisms to better understand the intersection of Plxna4, growth regulation and adiposity.

We observed striking behavioural responses to food in *plxna4* zebrafish mutants. Following food addition, mutants exhibited significantly increased distance travelled and average speed, indicating a heightened locomotor response compared to wild-type siblings. Mutants also showed greater speed variability, suggesting more erratic swimming patterns, which was further supported by an increased number of high-speed swim bursts. Interestingly, mutants also displayed more frequent resting periods, although this difference was not statistically significant. Altogether, these findings suggest that *plxna4* mutants are hyperactive in response to food, with increased movement irregularity compared to wild types. Prior studies in *Plxna4* mutant mice have also reported hyperactivity, though not specifically in response to food (39). These results raise intriguing questions about how these behavioural changes might relate to food consumption patterns, such as binge eating tendencies, or potential links to psychiatric traits. Considering PLXNAs role in neurogenesis, and recent research highlighting structural alterations - such as neuronal loss and reduced synaptic plasticity – that contribute to psychiatric disorders like anxiety and depression, disruptions to PLXNA4 could be associated with psychiatric behaviours (40). Future studies should examine feeding behaviours in greater detail and explore broader neuropsychiatric implications, particularly the relationship between Plxna4, hyperactivity, and food-driven behaviours.

## Acknowledgements

We thank Iris Pruñonosa Cervera, Susanna Riley, Aki Karas and Bronwyn Sell for initial *plxna4* mutant experiments and help developing the Ilastik segmentation workflow. We thank Bioresearch and Veterinary Services staff at the University of Edinburgh for general zebrafish husbandry, and Dr Carl Tucker and Paul Strachan for providing *Artemia* for feeding assays. Funding for the project was from the MRC (MR/S025685/1) and BBSRC (BB/X009467/1).

## Author contributions

PT and JENM conceived and designed the study, PT, ZL and JENM performed the experiments and analysis, PT, MC and JENM wrote and edited the manuscript, PT and JENM supervised the project.

## Data and code availability

Scripts used for analysis are available at https://github.com/jeminchin/plxna4_zebrafish

