## Supplemental Figures for "Human genetic studies and zebrafish models identify Plxna4 as a regulator of adiposity, somatic growth, and feeding behaviours"

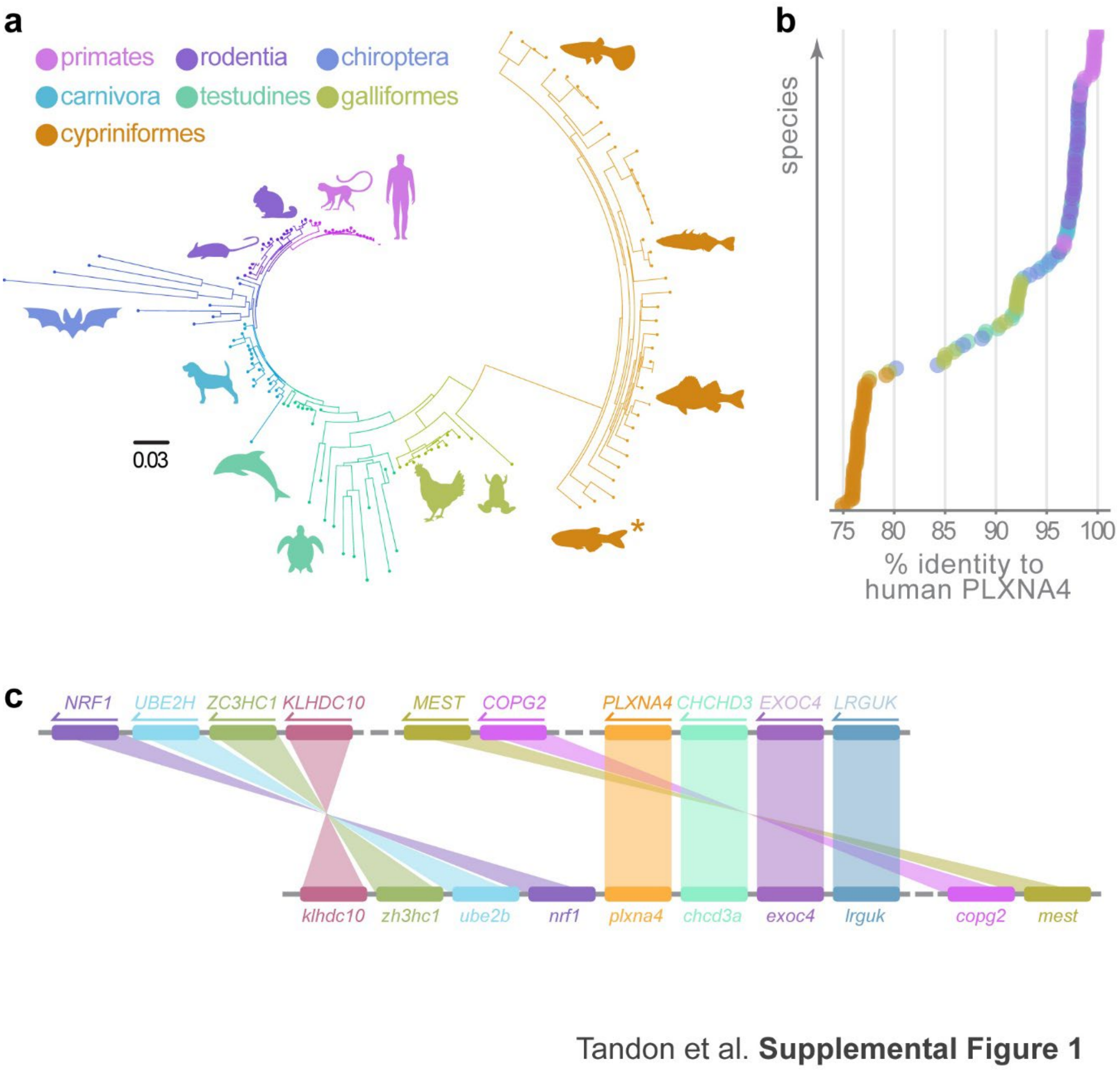


**Supplementary Figure 1. Sequence and syntenic conservation between zebrafish and mammalian Plxna4. (A)** Phylogentic tree showing relatedness for Plxna4 amino acid sequences across vertebrates. Sequences were obtained from Ensembl Genetree and the tree was generated using FigTree (https://github.com/rambaut/figtree/releases). **(B)** Percent amino acid identity for each species plotted relative to human PLXNA4. **(C)** Syntenic organisation of the human (upper row) and zebrafish (lower row) Plxna4 loci.

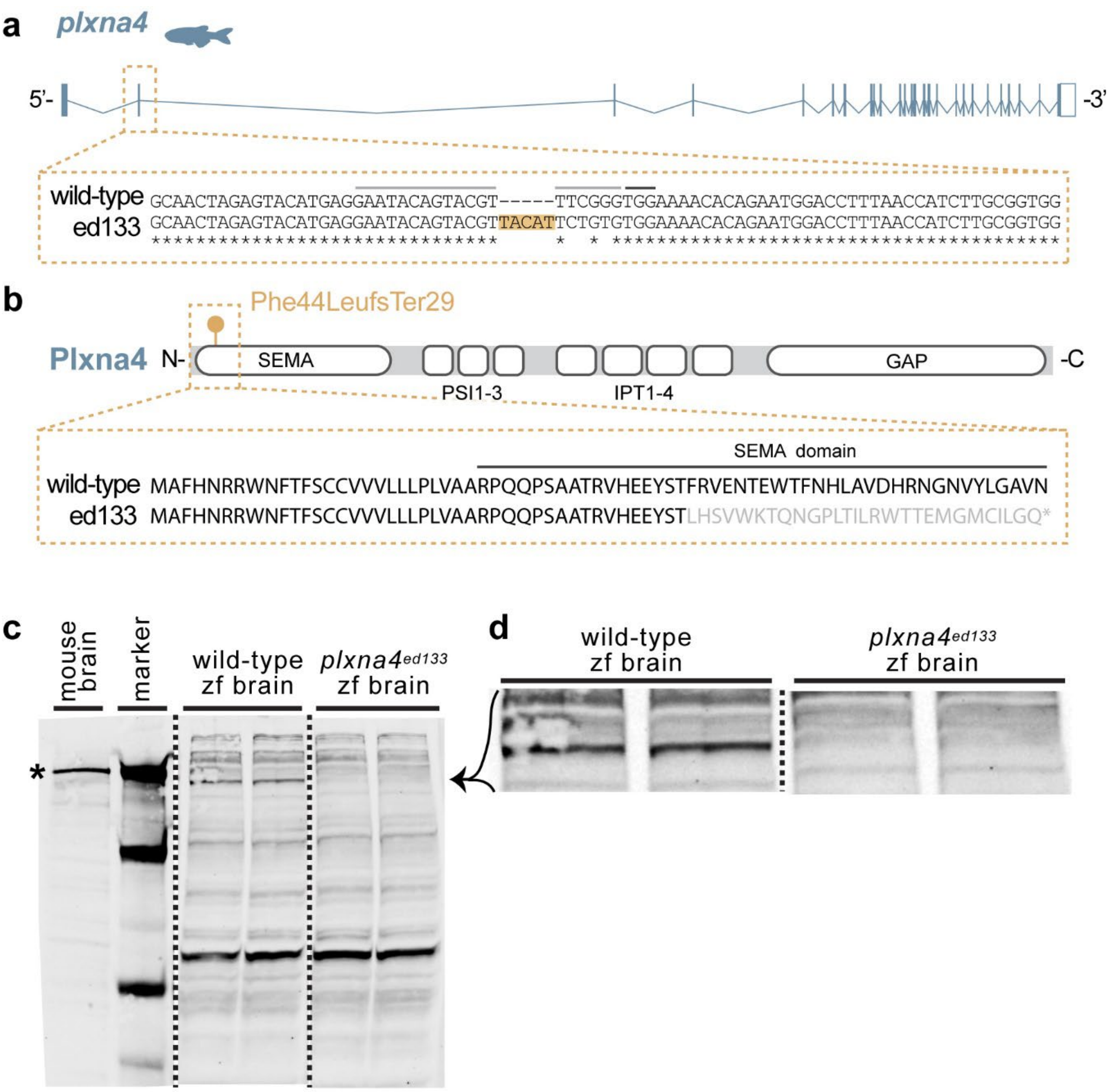


Tandon et al. **Supplemental Figure 2**

**Supplementary Figure 2. Generation of ed133 allele and verification of protein loss.** **(A,B)** Schematics of the ed133 allele) showing DNA and amino acid sequences (A and B respectively). **(C,D)** Full Western blot showing all bands from the anti-Plxna4 antibody. Note all bands are identical between wild-types and mutants except for a single band highlighted in D. The disappearing band in D is at the same molecular weight as mouse Plxna4 (marked by asterisk in C).

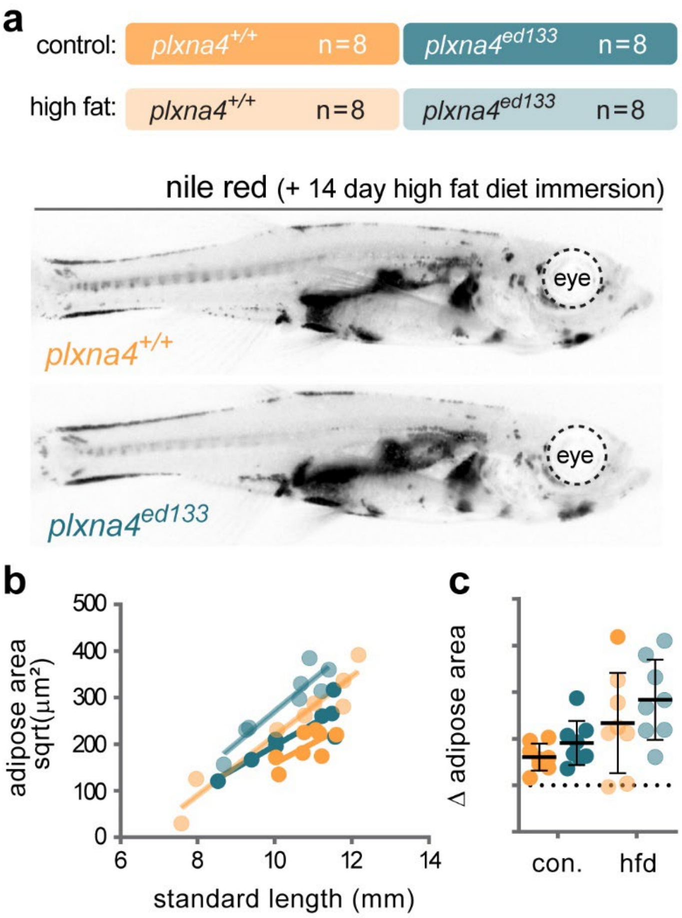


Tandon et al. **Supplemental Figure 3**

**Supplementary Figure 3. High-fat diet immersion experiment.** **(A)** Sample sizes for the four groups, and representative Nile Red images following 14-days of high-fat diet immersion. **(B)** Adipose areas following the 14-day diet manipulation. Only genotype and diet were statistically different by mixed-effects ANCOVA. Interaction between factors was not significant. **(C)** Change in adipose area after 14-days high fat diet. Note only genotype and diet were significant.bior
